# Membrane Lipids and Water Shape the Unbinding Kinetics of ZM241385 from Human Adenosine A_2A_ Receptor Variants

**DOI:** 10.64898/2026.09.07.749808

**Authors:** Marta Devodier, Davide Mandelli, Giulia Rossetti, Paolo Carloni

## Abstract

Ligand residence time is increasingly recognized as an important determinant of drug efficacy. In the human adenosine A_2A_ receptor (hA_2A_R) bound to the high-affinity antagonist ZMA, mutations at the extracellular entrance of the binding pocket markedly accelerate ZMA dissociation while largely preserving binding affinity. This effect has been attributed to disruption of the E169^ECL2^-H^7.29^ salt bridge that constrains the extracellular entrance in the wild-type receptor. However, how the interaction network is reorganized after salt-bridge disruption remains unknown. Here, we combine well-tempered metadynamics and infrequent metadynamics simulations to characterize the ZMA dissociation pathway and transition-state ensemble. The simulations reproduce the experimentally observed trend in dissociation rates and reveal that the hydrogen-bond network stabilizing ZMA in the bound state is largely replaced by water-mediated interactions during unbinding. In the mutants, disruption of the E169^ECL2^–H^7.29^ interaction expands the binding pocket and increases hydration, while additional ligand–lipid interactions emerge specifically at the transition state. These interactions are substantially less frequent in the wild type, indicating that membrane lipids can directly participate in ligand escape when the extracellular gate is disrupted. Thus, the mutations selectively reshape the transition-state interaction network – including recruitment of membrane lipids. Thus, the interaction of the latter with the ligand accompanies and may contribute to the mutants’ reduced dissociation barrier, thereby facilitating ligand release.

**Significance:** Drug efficacy depends not only on how tightly a ligand binds its target, but also on how long it remains bound. Why mutations can dramatically alter residence time without substantially changing binding affinity remains poorly understood. Here, we uncover a solvent-mediated mechanism of ligand dissociation from an important pharmaceutical target, the adenosine receptor A2A. For this system, the bound-state hydrogen-bond network is replaced by water-mediated interactions during unbinding, while lipid interactions emerge specifically in the receptor variants. These findings identify receptor–lipid interactions along the dissociation pathway as a potential target for rationally tuning drug kinetics.

## Introduction

Ligand residence times (RTs) have emerged as a key parameter in drug discovery, alongside binding affinities, because they may correlate with in vivo efficacy. Understanding the molecular determinants of ligand dissociation has therefore become a major objective in rational drug design (1–8).

In drug-design projects involving the optimization of RTs (9), a central unresolved question is why apparently modest changes in ligand binding can produce dramatic changes in ligand residence time. The human adenosine A2A receptor bound to the antagonist ZMA or ZM241385^1^ (Fig. 1A,B) offers an exceptional opportunity to address this problem. Mutations of residues located close to the ZMA binding site (E169^ECL2^→Q, T^6.58^ →A and H^7.29^→A, hereafter EtoQ, TtoA, HtoA, respectively) accelerate ZMA dissociation by more than an order of magnitude, while producing only modest changes in binding affinity (10) (see Table 1).

**Fig. 1.**
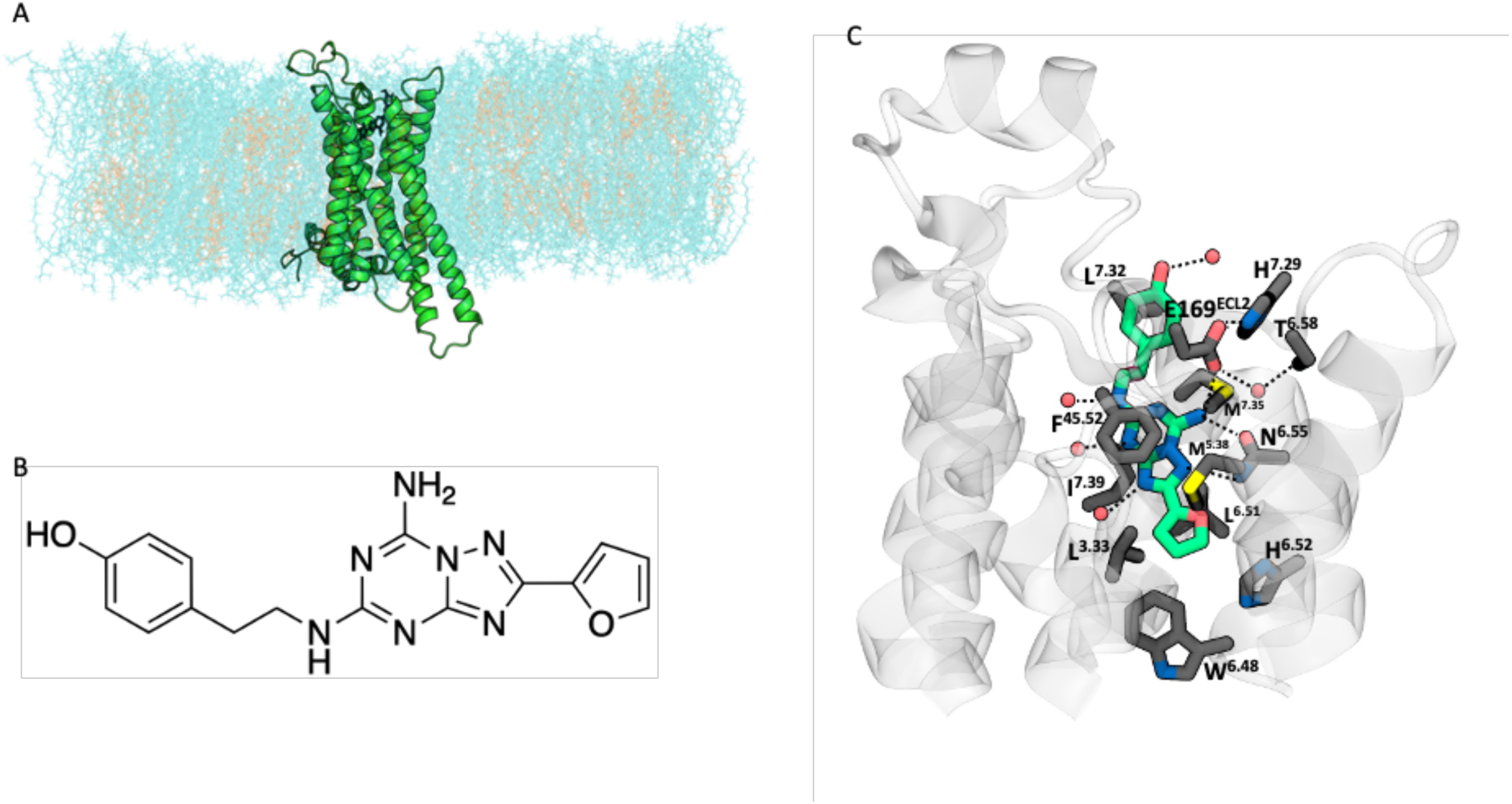
**(A)**: hA_2A_R (green ribbon) bound to ZMA (black) and embedded in a lipidic bilayer membrane (water not shown for clarity). (**B**) Chemical structure of ZMA (atom labeling in Fig. S1). **(C)**: X-ray structure of the hA_2A_R/ZMA complex (PDB ID:4EIY). Main residues interacting with the ligand (green licorice), as well as the E169^ECL2^-H^7.29^ salt bridge and T^6.58^ interacting with it, are highlighted. Hydrogen bonds between the ligand and the protein and water molecules are represented with dotted lines.

**Table 1.** For ZMA interacting with WT hA_2A_R and the mutants we report: (i) Residence time (RT [min]) and dissociation rate constant (k_off_ [min^-1^]) from experiment (10) and from InMetaD; (ii) p-values (p) used to test the statistical confidence of the computed k_off_ values (details in Fig. S2); (iii) binding free energy (ΔG_0_ [kcal/mol]) from experiment (10) and from WT-MetaD; (iii) predicted free energy barriers to ligand unbinding from WT-MetaD (ΔG^‡,sim^ [kcal/mol]). No results are provided for the predicted ΔG_0_ and ΔG^‡,sim^ of HtoA, as the WT-MetaD simulation could not converge.

| | $RT^{\text{exp}}$ | $RT^{\text{sim}}$ | $k_{\text{off}}^{\text{exp}}$ | $k_{\text{off}}^{\text{sim}}$ | $p$ | $\Delta G_0^{\text{exp}}$ | $\Delta G_0^{\text{sim}}$ | $\Delta G^{\ddagger, \text{sim}}$ |
| --- | --- | --- | --- | --- | --- | --- | --- | --- |
| **WT** | 84.0 | 480.7 | 0.012 | 0.002 | 0.61 | -11.1 | -9.5 | 7.6 |
| **EtoQ** | 1.4 | 5.9 | 0.733 | 0.169 | 0.30 | -10.6 | -8.3 | 5.8 |
| **TtoA** | 5 | 0.45 | 0.199 | 2.218 | 0.69 | -10.5 | -8.6 | 6.9 |
| **HtoA** | 4.5 | 6.2 | 0.224 | 0.161 | 0.15 | -10.9 | — | — |

This striking difference between thermodynamics and kinetics suggests that the mutations likely stabilize the bound state similarly and modulate the free-energy barrier governing ligand escape. Previous structural and computational studies provide important clues to the molecular origin of this effect. In the bound state, ZMA is stabilized by an extensive network of direct protein-ligand and water-mediated interactions within the orthosteric binding pocket (11) (Fig. 1C). Moreover, enhanced-sampling simulations previously identified the extracellular loops as important determinants of ZMA recognition and highlighted the E169^ECL2^–H^7.29^ salt bridge – in which H^7.29^ is double-protonated – as an “access-control” interaction at the extracellular entrance of the binding pocket (12). These simulations suggested that remodeling of this extracellular interaction network, together with the surrounding membrane environment, can influence ligand access to the orthosteric site. More recently, NMR experiments (13) showed that the disruption of the E169^ECL2^–H^7.29^ salt bridge at the entrance of the binding pocket (Fig. 1C) caused by the EtoQ, TtoA and HtoA mutations correlates with residence time, providing experimental evidence of mutation-induced remodeling of this extracellular gate. Taken together, these observations suggest that disruption of the E169^ECL2^–H^7.29^ interaction may facilitate ZMA escape by altering the structural and dynamic environment encountered along the dissociation pathway, thereby lowering the corresponding free-energy barrier and reducing ligand residence time, as experimentally observed (Table 1).

Yet these observations leave a fundamental question unanswered: what interactions replace the ligand–salt-bridge interactions as the ligand crosses the dissociation barrier after the salt bridge has broken? Experimental techniques, including high-resolution X-ray crystallography, cryo-EM, and NMR spectroscopy, provide invaluable information on bound-state structures and conformational changes, but they cannot directly characterize the structural determinants of the transient transition-state ensemble (TSE) governing the dissociation free-energy barrier. Enhanced-sampling molecular simulations can characterize unbinding pathways and thereby provide an atomistic description of the TSE, while also enabling a qualitative estimation of residence times (RTs) (14). Here, we address this question by combining infrequent (In-) and well-tempered (WT-) metadynamics (MetaD) (15,16). Our aim is to reconstruct the ZMA unbinding pathway and characterize the interaction network stabilizing the TSE. Our predictions are in good agreement with the k_off_ trend (Table 1). A detailed analysis of the unbinding trajectories reveals how the sizable protein-ligand hydrogen-bond network observed in the bound state is mostly replaced at the TS by hydrogen bonds with water and, for the variants, with lipids. Furthermore, we show that hydrophobic interactions constitute the principal persistent protein–ligand contacts during the unbinding process. These findings reveal a previously unrecognized solvent-mediated mechanism underlying ligand dissociation from hA_2A_R and suggest that mutation-specific interactions with lipids contribute to the profound changes in residence time observed for the variants, without equivalently affecting the binding affinity.

## Results

Table 1 compares the predicted and experimental k_off_ values, obtained by In-MetaD simulations. The computed k_off_ values reproduce the experimental trend. Their accuracy is in line with the current state-of-the-art techniques (14).

The binding free energies of WT, EtoQ, TtoA hA_2A_R were calculated from WT-MetaD simulations. They feature a statistical error of 1 kcal/mol, as observed experimentally, and deviate from the experimental values by ∼2 kcal/mol, consistent with the accuracy typically reported for ligand binding to GPCRs (17,18). The values for HtoA mutant are not reported as the WT-MetaD did not converge and hence its results are not reported here.

Discrepancies between simulations and experiments can be caused, at least in part, by force-field inaccuracies (19,20) as well as differences between the experimental and computational setups.

To gain structural insight into the molecular determinants underlying the computed kinetic and thermodynamic values, we investigate the free energy landscapes as a function of the ligand position *z* along the unbinding axes and the ligand RMSD with respect to its pose in the binding site (RMSD_ZMA_ hereafter) (Fig. 2). Because the WT-MetaD free energy landscape of HtoA did not converge, HtoA was excluded from this analysis.

**Fig. 2.**
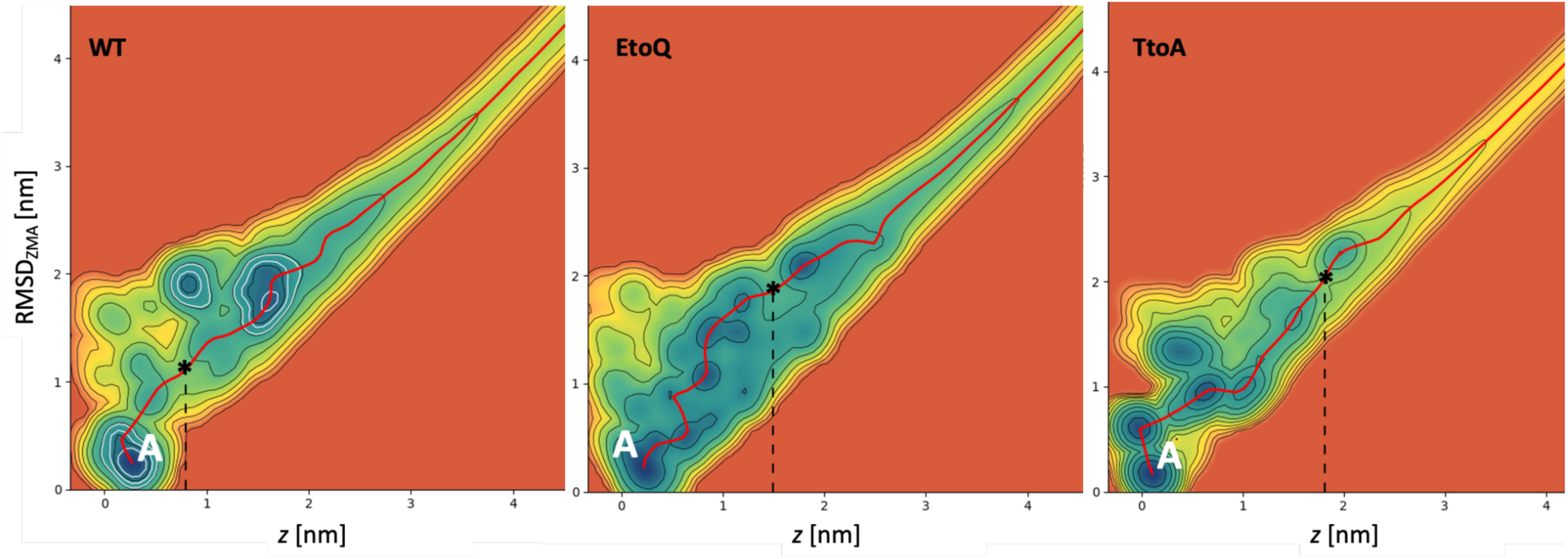
Free energy landscapes of (a) WT hA_2A_R, (b) EtoQ and (c) TtoA mutants. Stars mark the transition state (TS) along the minimum free energy path (red line). Dashed vertical lines drop from each star to the x-axis, indicating the corresponding *z* value. Free energies were obtained by reweighting the WT-MetaD trajectories, following the method of Ref. (21).

In WT hA_2A_R, basin A corresponds to the global free energy minimum. In this structural ensemble, the ligand is accommodated in its binding site. The receptor cavity volume is 0.34 ± 0.04 nm^3^, and it is highly hydrated, containing on average 30±4 water molecules (Table S1 in SI). The ZMA pose closely resembles the X-ray pose (11) (0.1 nm <RMSD_ZMA_ < 0.3 nm, see Fig. 3A). The ligand maintains its hydrophobic contacts with nonpolar groups of 12 residues (Table S2 in SI), it forms *π*–*π* stacking interaction with F^45.52^, along with three H-bonds with the protein (ZMA_N4_ - E169^ECL2^_OE_, ZMA_N4_ - N^6.55^_OD_, ZMA_N4_ - N^6.55^_ND_) and four H-bonds with water molecules (see Figs. 1C, 3A and 4). The maxima of the H-bond length distributions almost overlap with the corresponding distance that can be inferred by X-ray values (Fig. 4). The E169^ECL2^–H^7.29^ salt bridge, which acts as a barrier for ZMA dissociation, is largely maintained (∼40% of the frames, comparable with the percentage in the unbiased simulation of the bound state (Section S1 of SI).

**Fig. 3.**
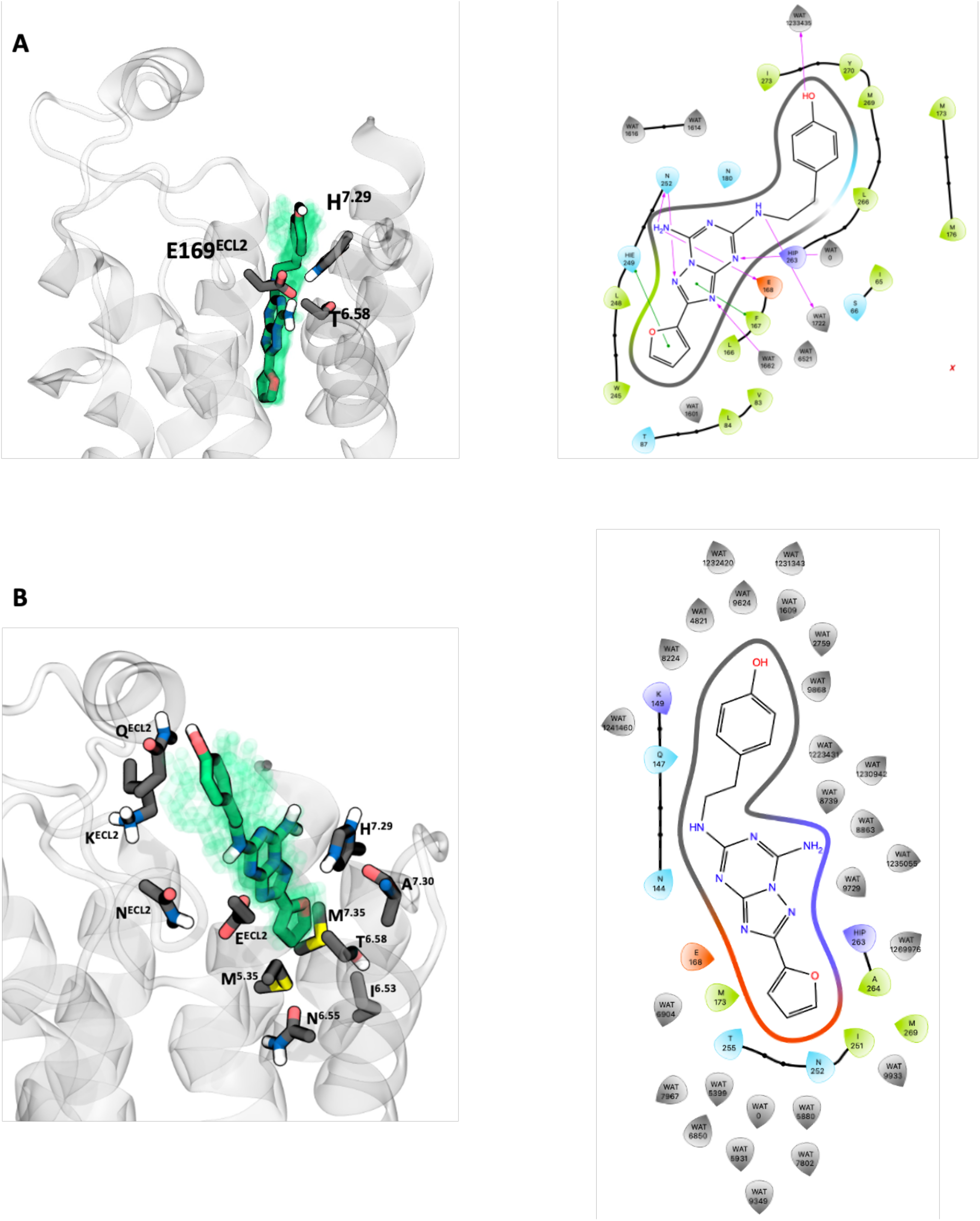
Representative structures of (A) the bound state and (B) transition state of WT hA_2A_R obtained in WT-MetaD simulations. (A, left panel) An ensemble of structures (within 1 kBT from the minimum) is represented in shaded green, while the crystallographic pose (11) is shown in licorice representation. The salt-bridge and T^6.58^ interacting with it are also shown. The corresponding 2D scheme (right panel) shows the interactions of the ligand within the binding pocket. (B) The same analysis is reported for the transition state. The ensemble of structures has been extracted considering a region of 1kBT around the saddle point, and the 2D scheme (right panel) shows the residues and water molecules interacting with the ligand.

**Fig. 4.**
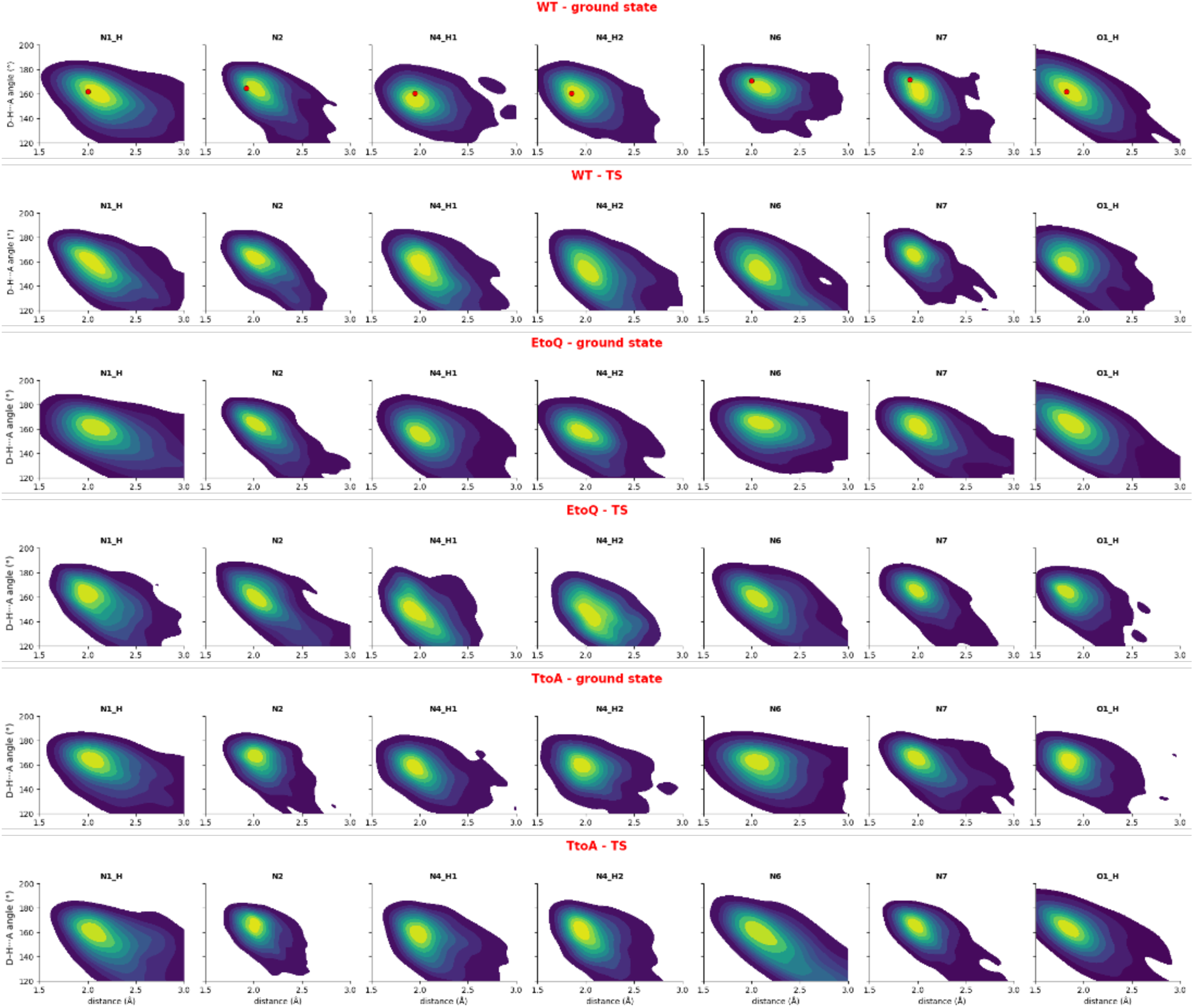
WT-MetaD simulations. Distributions of ZMA H-bond distances and angles with water molecules and protein in the WT hA_2A_R, and EtoQ and TtoA mutants in basin A and at the TS. Atoms of the ligand are labeled according to the definition in Fig. S1 of the SI. Red dots mark the hydrogen bond geometry inferred from the X-Ray structure (PDB ID:4EIY). Full distributions are reported in Figs. S3-S8 of the SI.

Ligand dissociation along the minimum free energy paths (MFEP, red line in Fig. 2) to the TS (indicated by a star in Fig. 2) is associated with a free-energy barrier of about 8 kcal/mol. In the TSE, the E169^ECL2^_OE_ - ZMA_N4_, N^6.55^ - ZMA_N4_ and N^6.55^ - ZMA_N4_ H-bonds are lost. ZMA occasionally forms weak hydrogen bonds with E^ECL2^ and H^7.29^. It also forms as many as five hydrogen bonds with water molecules (Figs. 3B, 4). Only two hydrophobic contacts with M^5.35^ and M^7.35^ are present, while the salt bridge is completely disrupted (an average distance of 0.847 nm ± 0.014). The ligand forms few interactions with lipids (24% of the frames) and it is fully hydrated (Table S3 of SI): 21±3 water molecules are present within 0.35 nm of the heavy atoms of the ligand.

We next focus on the EtoQ hA_2A_R mutant. In the absolute free energy minimum (A) the ligand’s pose is similar to that of the WT protein: the ligand maintains its hydrophobic contacts with nonpolar groups of 12 residues (Table S2 of SI) and *π*–*π* stacking interaction with F^45.52^, along with two out of three H-bonds with the protein (the ZMA_N4_ - E169^ECL2^_OE_ interaction in the WT is here replaced by either Q169_OE_ - or water-ZMA interactions) and four H-bonds with water molecules (see Figs. 4, 5A). The cavity volume (0.40 ± 0.06 nm^3^) is larger than in the WT and consequently contains on average more water molecules (35±4) (Table S1 of SI).

**Fig. 5.**
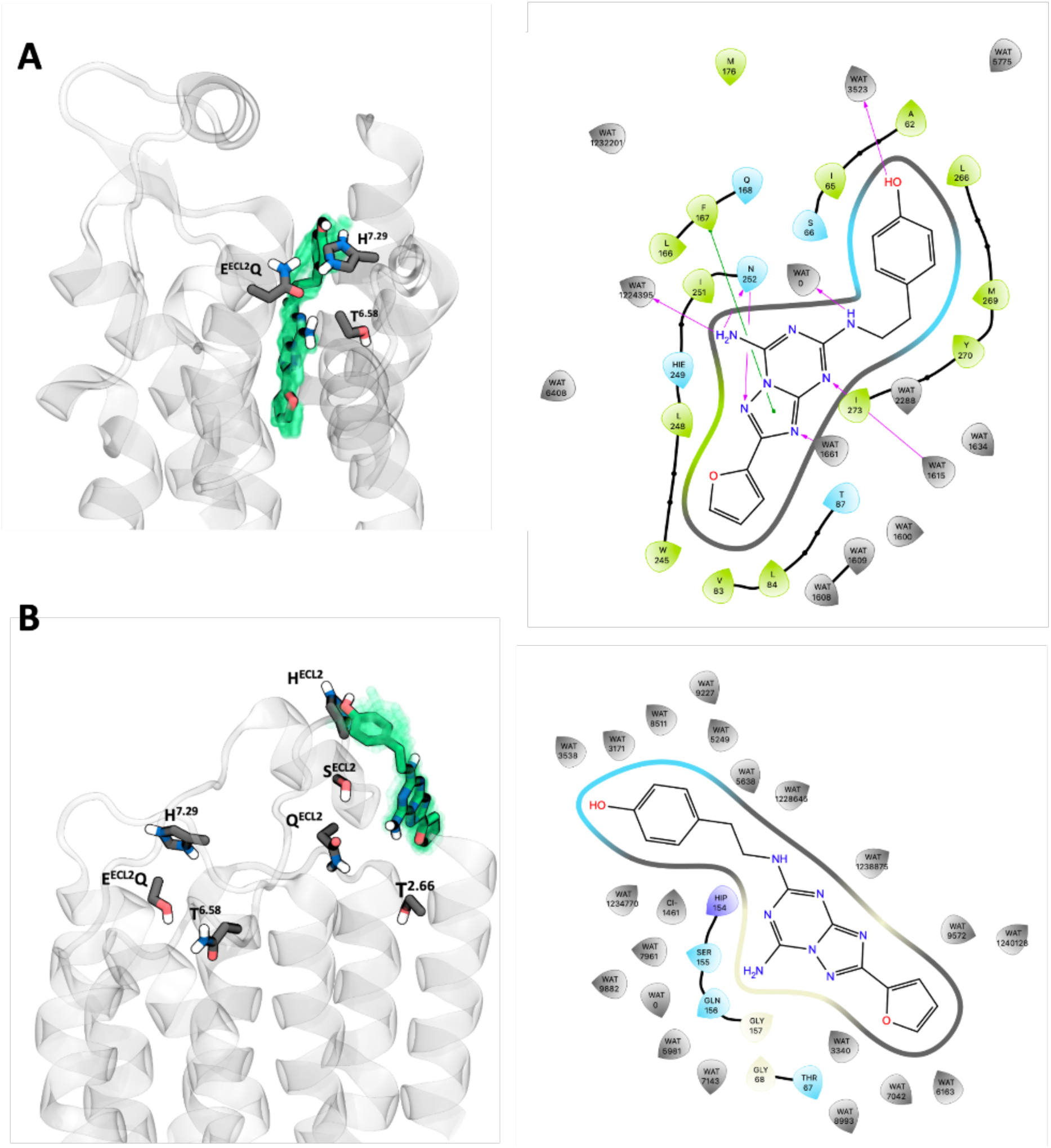
Representative structures of (A) the bound state and (B) transition state of EtoQ, depicted as in Fig. 3.

Ligand dissociation along the MFEP to the TS is associated with a free-energy barrier of about 6 kcal/mol. The ligand loses its hydrogen bonds with N^6.55^ at ZMA and N^6.55^ at ZMA (Fig. 5B) but occasionally forms two new ones with the protein (specifically, between ZMA_N4_ and T^2.66^ and S155^ECL2^; Figs. 4 and 5B). It also forms as many as five hydrogen bonds with water molecules (Figs. 4, 5B). In contrast to the WT protein, the ligand forms persistent contacts with lipids (98% of the frames). The residues Q169 and H^7.29^ do not interact with the ligand. In the TS, the ligand is fully hydrated as in the WT (Table S3 of SI).

In the global free energy minimum A of TtoA hA_2A_R, the ZMA pose is similar to that of the WT (Fig.6A): the ligand maintains its hydrophobic contacts with 11 residues (Table S2 of SI) and *π*–*π* stacking interaction with F^45.52^, along with two out of three H-bonds with the protein^2^ and four H-bonds with water molecules (Fig. 6A). The cavity volume (0.37±0.03 nm^3^) is intermediate between that of WT and that of EtoQ, containing 36 ± 5 water molecules (Table S1 of SI).

**Fig. 6.**
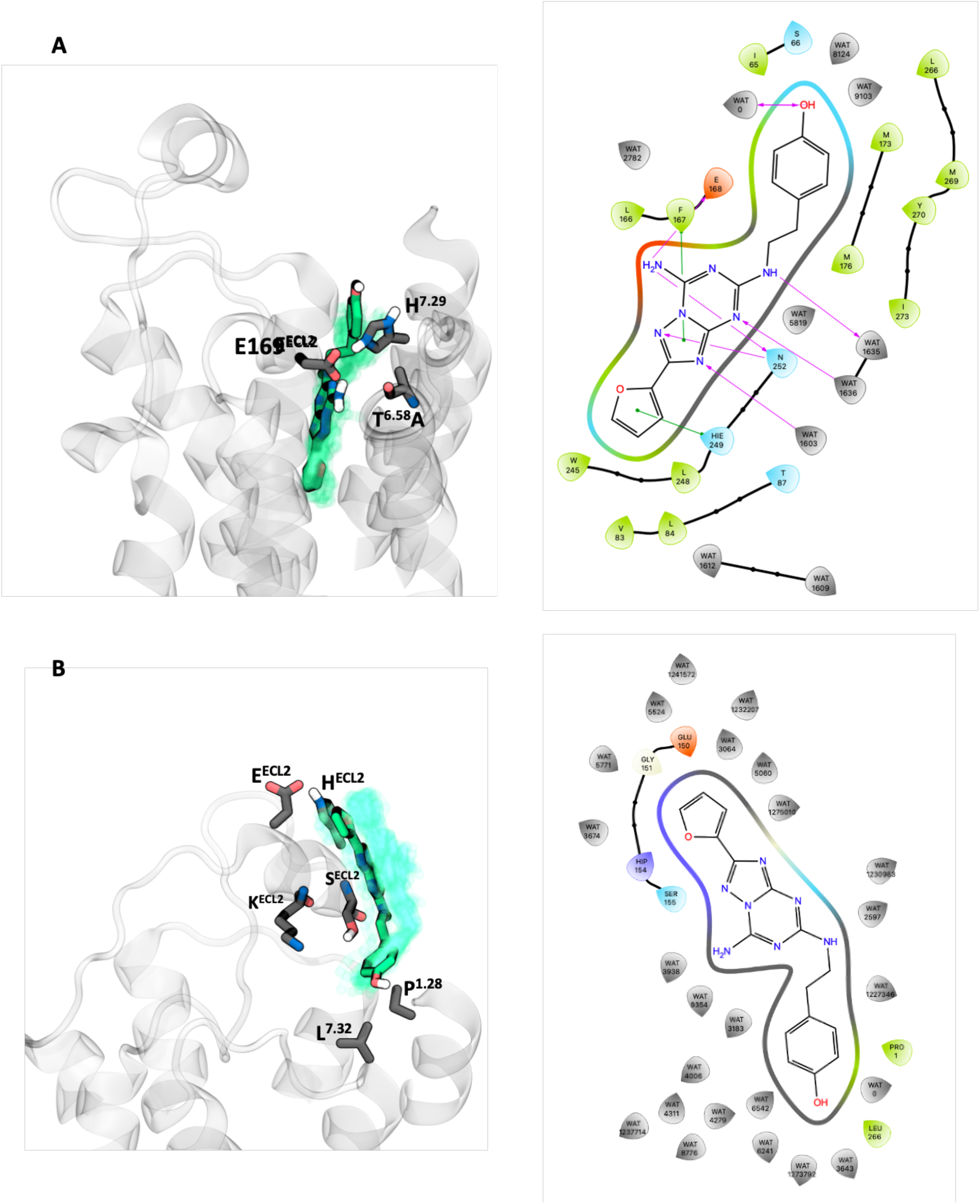
Representative structures of (A) the bound state and (B) transition state of TtoA, depicted as in Fig. 3.

Ligand dissociation along the MFEP to TS is associated with a free-energy barrier of about 7 kcal/mol. The ligand loses all its hydrogen bonds with the protein except an H-bond with S155^ECL2^, replaced at times by water and forms as many as six hydrogen bonds with water molecules (Figs. 4, 6B). Two hydrophobic contacts are present with P^1.28^ and L^7.32^. As in the previous mutant, the ligand forms more contacts with lipids (52% of frames) than the WT. The residues E169^ECL2^ and H^7.29^ do not interact with the ligand. The TS is fully hydrated as in the WT protein, with 20±3 water molecules present within 0.35 nm of the heavy atoms of the ligand (Table S3 of SI).

### Comparison between the WT and the mutants

The dissociation free energy of the WT protein, as obtained from the MFEP analysis, is 1-2 kcal larger than those of the mutants, consistently with the predicted difference in RTs (Table 1). The protein forms contacts with the ligand at atoms N4 and N6 – which are within the dominant protein-anchoring interactions in the bound state (Fig. 1) – but they are basically lost at the TS in WT, EtoQ and TtoA (Figs. 3B,5B,6B). In EtoQ, instead, N4 still occasionally forms a hydrogen bond. Overall, we conclude that the transition state is then characterized by a loss of specific protein hydrogen bonds, with solvent occupying these positions instead.

## Discussion

Our simulations reproduce the experimentally observed trend in RTs with the expected accuracy for this type of calculations, while the calculated binding free energies remain similar among the systems within the statistical uncertainty of the calculations. The WT receptor exhibits a barrier of approximately 8 kcal/mol, compared with approximately 6 and 7 kcal/mol for EtoQ and TtoA, respectively. Although small, a reduction of approximately 1 kcal/mol is sufficient to produce a very large change in the dissociation rate due to the exponential sensitivity of kinetic rates to the free-energy barrier, and is in line with the observed dramatic decrease in RT for the mutants.

Together, these results provide a molecular explanation regarding the fact that the mutations primarily affect the kinetic barrier to dissociation rather than the thermodynamic stability of the bound state. In the WT receptor, ZMA is stabilized in the binding pocket by a combination of hydrogen bonds and hydrophobic contacts, while the E169^ECL2^–H^7.29^ interaction at the extracellular entrance is a barrier for unbinding. In contrast, the EtoQ and TtoA mutations destabilize this interaction, as already inferred by NMR (13). As ZMA proceeds from the bound state toward the transition state, the hydrogen-bond network observed in the binding pocket is extensively reorganized. Three protein hydrogen bonds and four ligand–water hydrogen bonds in the bound state are replaced at the transition state by fewer direct protein interactions and a predominantly solvent-mediated interaction network. Importantly, hydrophobic contacts remain the principal persistent direct interactions between ZMA and the receptor. Thus, ligand escape does not correspond simply to the progressive loss of all protein contacts but rather involves a qualitative transition from a specific protein–ligand H-bond interaction network to a more dynamic, solvent-mediated environment. This is fully consistent with previous studies (18,19) that suggested that protein–ligand interactions are transiently replaced by solvent-mediated contacts.

The EtoQ and TtoA mutations also lead to an increase in binding pocket volume and water content, while at the transition state, the ligand is highly hydrated across all systems (Tables S1, S3 of SI). In addition to water-mediated interactions, ZMA forms several contacts with lipids at the TS, in contrast to the WT receptor. To our knowledge, such ligand–lipid interactions at the transition state have not been reported previously. This finding indicates that, as the ligand approaches the TS and the extracellular binding pocket opens, interactions with the surrounding membrane can become part of the interaction network encountered during ligand dissociation. Thus, the membrane may participate not only in the initial recognition and vestibular stabilization of ZMA, as previously proposed (12), but also in shaping the interaction network at the transition state.

Our results also suggest that solvent should be considered an active component of the dissociation mechanism rather than merely the background medium in which ligand unbinding occurs. The replacement of relatively rigid protein–ligand hydrogen bonds by dynamic water-mediated interactions may increase the configurational freedom of both the ligand and its surrounding solvent. Such solvent reorganization could therefore contribute to the free-energy barrier and help determine the kinetics of escape. Thus, solvent reorganization is a structural feature that accompanies and potentially contributes to the reduced dissociation barriers of the mutants.

### Conclusions

We have presented a molecular simulation study on the kinetics of unbinding of the ZMA ligand from its target hA2A receptor and mutants with accelerated dissociation and shorter residence times. As the ligand approaches the transition state, specific protein–ligand hydrogen bonds are largely replaced by water-mediated interactions: the dominant protein-anchoring interactions in the bound state at atoms N4 and N6 (Fig. 1C) are either completely (in WT and TtoA) or almost completely (in EtoQ) lost in the transition state (Figs. 3B,5B,6B). The latter is characterized by a loss of specific protein hydrogen bonds, with solvent (and lipids for the protein mutants) occupying these positions instead. This dynamic remodeling of protein, ligand, lipids and solvent provides a molecular framework for understanding how receptor mutations can strongly alter residence time while leaving binding affinity comparatively unchanged.

The present results should not be interpreted as demonstrating that solvent-mediated stabilization is a universal mechanism for GPCR ligand dissociation. The analysis concerns one ligand, one receptor, and three protein species, and the calculated kinetic and thermodynamic quantities retain the uncertainties inherent to enhanced-sampling simulations and the underlying force field. Nevertheless, the consistency between the simulated and experimental dissociation trends, together with the common transition-state hydration pattern observed across the systems, provides strong evidence that solvent reorganization is an important component of the ZMA dissociation mechanism, as seen for other systems. The newly observed lipid interactions at the transition state further indicate that the membrane environment may participate in the dissociation process.

## Materials and Methods

### System Setup and MD simulations WT hA_2A_R

The crystal structure of the WT hA_2A_R in complex with its high-affinity antagonist ZMA bound within the OBS was taken from the PDB (ID: 4EIY). All expression tags and the apocytochrome b562RIL inserted in the ICL3 loop were removed to restore the WT hA_2A_R sequence. Missing loops and residues were modelled with MODELLER (22) and the system was then embedded in a heterogeneous bilayer system using the CHARMM-GUI server (23,24). The membrane consists of three different lipids with the following composition: cholesterol (30%), POPC (50%), POPE (20%). Finally, water and Na^+^ and Cl^-^ ions were added to solvate and neutralize the system at an ionic strength of 150 mM. The final WT system consists of 231,639 atoms in an orthorhombic box of size ∼(116 ⨉ 116 ⨉ 147) Å^3^.

The AMBER ff19SB force field (25), the Lipid21 force field (26), and the OPC model (27) were used for the protein, lipids, and water molecules, respectively. The General Amber force field (GAFF) (28) parameters were used for ZMA, along with RESP (29) atomic charges using the Gaussian 09 code (30) with the HF-6-31G* basis set. Electrostatic interactions were treated with the Particle Mesh Ewald (PME) method (31) and the cutoff for nonbonded interactions was set to 12 Å. All covalent bonds involving hydrogen atoms were constrained using the LINCS algorithm (32). The system geometry was first minimized using the steepest descent algorithm, followed by a gradual heating phase in 40 steps to 310 K in 1 ns of annealing, in the presence of the restraints. The system was then successively equilibrated following a multiple-step protocol, with progressively decreasing harmonic restraints on protein backbone atoms, lipids and ligand. The first steps consisted of 20 ns NVT simulation (T = 310 K), in which the positional restraints on the protein, ligand and lipids were gradually released; the second steps consisted of 30 ns simulations in the NPT ensemble (T = 310 K, P = 1 bar), gradually releasing the restraints on the protein. In all these simulations, the selected timestep was 2 fs, and the temperature was controlled using the Bussi-Parrinello stochastic velocity rescaling thermostat (33) with a time constant of 0.1 ps. The thermostat was applied separately to the protein-ligand system, the lipid bilayer, and the solvent. Pressure was controlled using the Berendsen barostat (34) with a time constant of 1 ps. After equilibration, a 500 ns-long production run was carried out without restraints. During the production run, the temperature and the pressure (T = 310 K, P = 1 bar) were controlled, respectively, using the Nosé-Hoover thermostat (35) with a time constant of 0.5 ps, and the semi-isotropic \ Parrinello-Rahman barostat (36) with a time constant of 2 ps and a compressibility of 4.5⨉10^−5^ bar^−1^. All the other settings were the same as used during equilibration. All simulations were carried out with GROMACS v2024.3 (37). The final snapshot of the production run was used as the starting configuration for the enhanced sampling simulations, as well as to prepare the mutants.

### Mutants

Starting from the NPT equilibrated WT system, the EtoQ, HtoA and TtoA mutant models were obtained using UCSF Chimera (38), employing the Rotamers tool to select rotamers with minimal steric clashes. Each system was then equilibrated following the same procedure as for the WT model. The final NPT equilibrated structures were used as starting configurations for the enhanced sampling simulations.

### Enhanced-sampling simulations

For the WT system, well-tempered metadynamics (WT-MetaD) (16) was used to investigate the ligand unbinding process, while for the mutants we used multiple walkers WT-MetaD (39) with 6 walkers per system, following the approach used in Ref (17). In both cases, we restrained the center of mass (COM) of the ligand within a funnel-shaped volume (Fig. S9), following the setup of Ref. (40). Defining d as the vector connecting the COM of ZMA to the COM of the Cα atoms of the hA_2A_R binding-pocket residues, the chosen collective variables (CVs) are its component *z* along the funnel axis, and the radial distance *r* of the ligand COM from the axis (see Fig. S9). The deposition rate of the Gaussian kernels was set to 2 ps and the initial Gaussian height to 2 kJ/mol, with a bias factor of 15 and a Gaussian width of 0.05 nm for both CVs. Binding free energies were computed using Eq. 29 of Ref. (41) (Fig. S10), using *z* as the reaction coordinate, defining the bound state as 0.2 nm < z < 0.4 nm and the unbound state as 5.5 nm < z < 5.6 nm. We ensured that in the bound region there are no interactions of the ligand with the restraint. The volumetric correction is -k_B_T log(ℓΣC_0_), with ℓ=0.1 nm, Σ=πR^2^, where R=0.15 nm is the radius of the cylindrical part of the restraint, and C_0_=1 mol/l = 0.6022 nm^-3^ is the standard concentration.

Infrequent metadynamics (15) was used to investigate the structural and kinetic determinants of ligand dissociation using *z* as the CV. In practice, for each system, we ran 15 independent simulations starting with the ligand in the bound pose. The deposition rate of the Gaussian kernels was set to 50 ps and the initial Gaussian height to 0.6 kJ/mol, with a bias factor of 15 and a Gaussian width of 0.05 nm. The simulations were stopped once the ligand reached the unbound state (z > 4 nm). The characteristic unbinding time was obtained as the average of the rescaled times obtained from the 15 independent runs. A Kolmogorov-Smirnov (KS) test (42) was used to quantitatively assess the quality of the predictions.

All simulations were performed using GROMACS v2024.3 patched with PLUMED 2.9.3 (43).

## Author Contributions

MD performed all calculations, conducted all analyses and contributed to manuscript writing. DM, GR, and PC designed the project and contributed to manuscript writing.

## Acknowledgments

The authors acknowledge support by the European Union’s HORIZON MSCA Doctoral Networks program, under Grant Agreement No. 101072344, project AQTIVATE (Advanced computing, QuanTum algorIthms and datadriVen Approaches for Science, Technology, and Engineering). The authors gratefully acknowledge computing time on the JUWELS Cluster and Booster module of the Juelich Supercomputing Center (JSC) (44) under grant project *mopkin*.

## Supporting information for

**S1. How the mutations destabilize the salt-bridge interaction**

The WT, EtoQ, and TtoA variants underwent 500 ns of unbiased MD simulations. We analyzed the persistence of the salt bridge during the MD simulations by computing the minimum inter-atomic distance between E169^ECL2^_OE1/OE2_ - H^7.29^_ND1/NE2_ in the WT hA_2A_R and TtoA mutant, and between Q169_OE1_ - H^7.29^_ND1/NE2_ in the EtoQ mutant (Fig. S11). The results show that the interaction is persistent in the WT (∼ 50% of frames). In contrast, in both the EtoQ and TtoA mutants the frequency of the interaction is significantly reduced to ∼19% and ∼37% of frames, respectively.

In particular, mutation of the E169^ECL2^ residue leads to disruption of the salt bridge interaction, as indicated by the peak of the distribution moving to ∼0.5 nm, compared to ∼0.3 nm in the WT. On the other hand, mutation of T^6.58^ presents a bimodal distribution with the first peak at ∼0.3 nm and a second peak at ∼0.6 nm, indicating destabilization of the E169^ECL2^-T^6.58^-H^7.29^ triad and hence reduced persistence of the salt bridge.

**Fig. S1.**
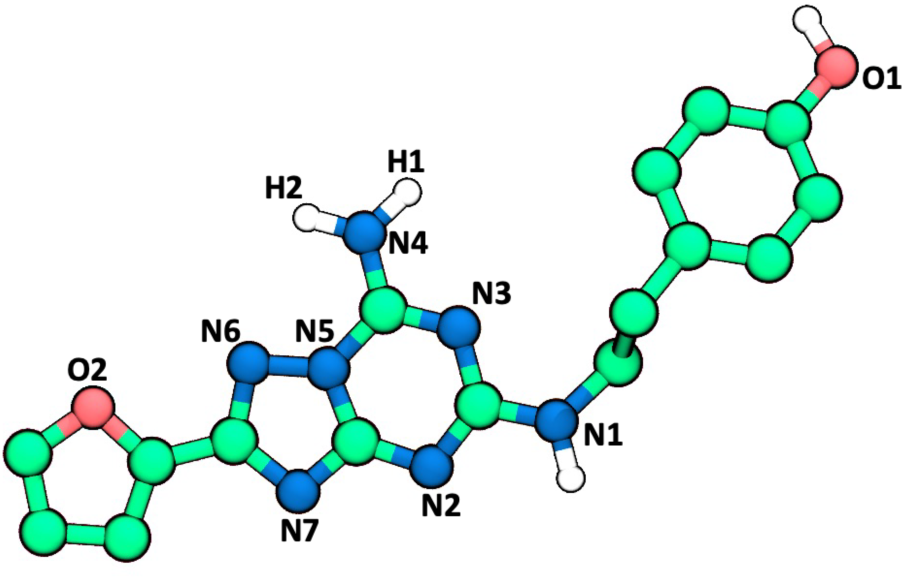
Enumeration of ZMA atoms as used in the main text.

**Fig. S2.**
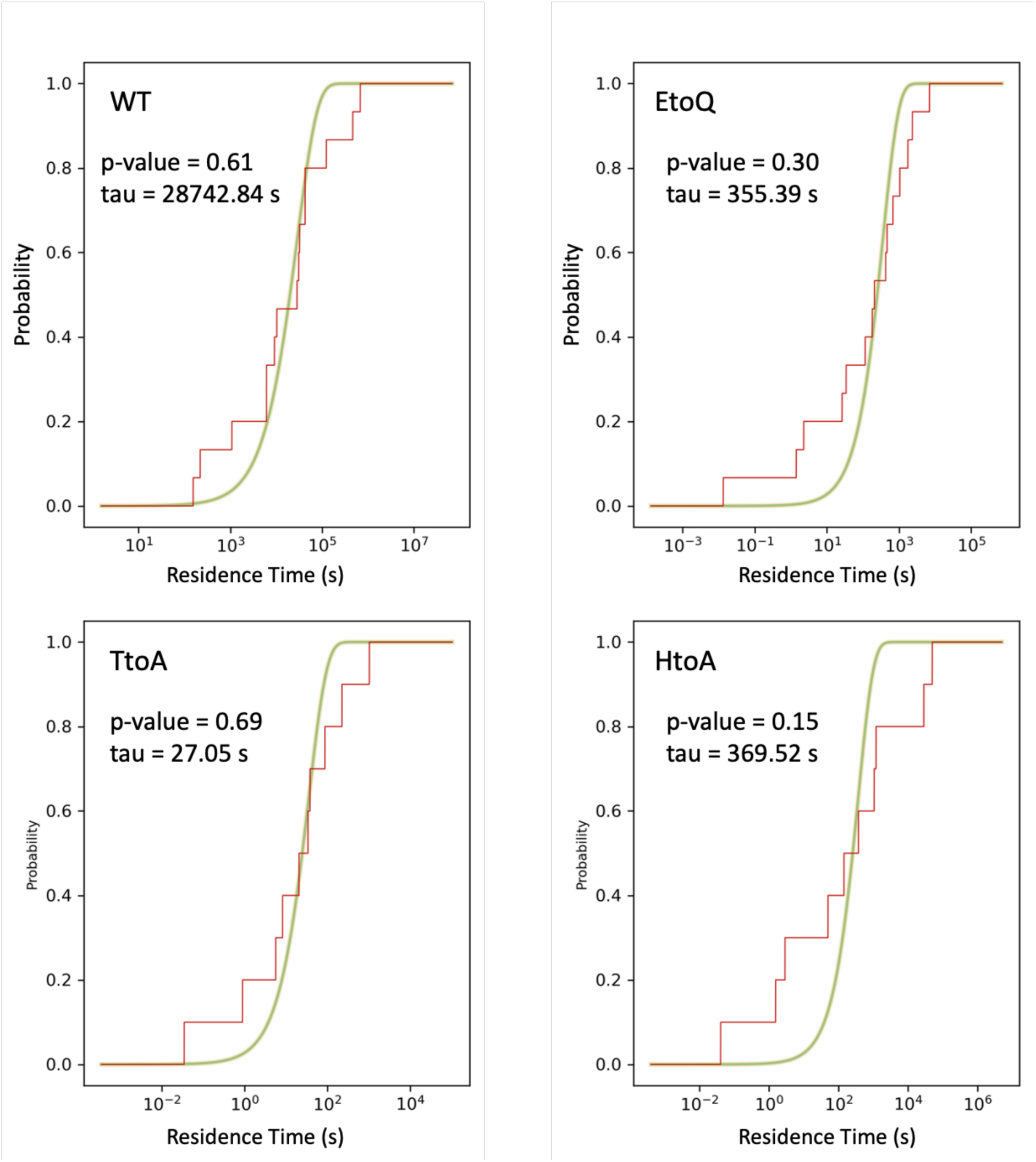
The cumulative distribution of escape times computed from infrequent metadynamics simulations on the WT hA_2A_R, EtoQ, HtoA and TtoA mutants. The labels indicate the ligand’s RT, and the tau value obtained from a fit of the curve, and the p-value used to test the statistical confidence of the fit. The computational results reproduce well the experimental trend (see Table 1 in the main text).

**Fig. S3.**
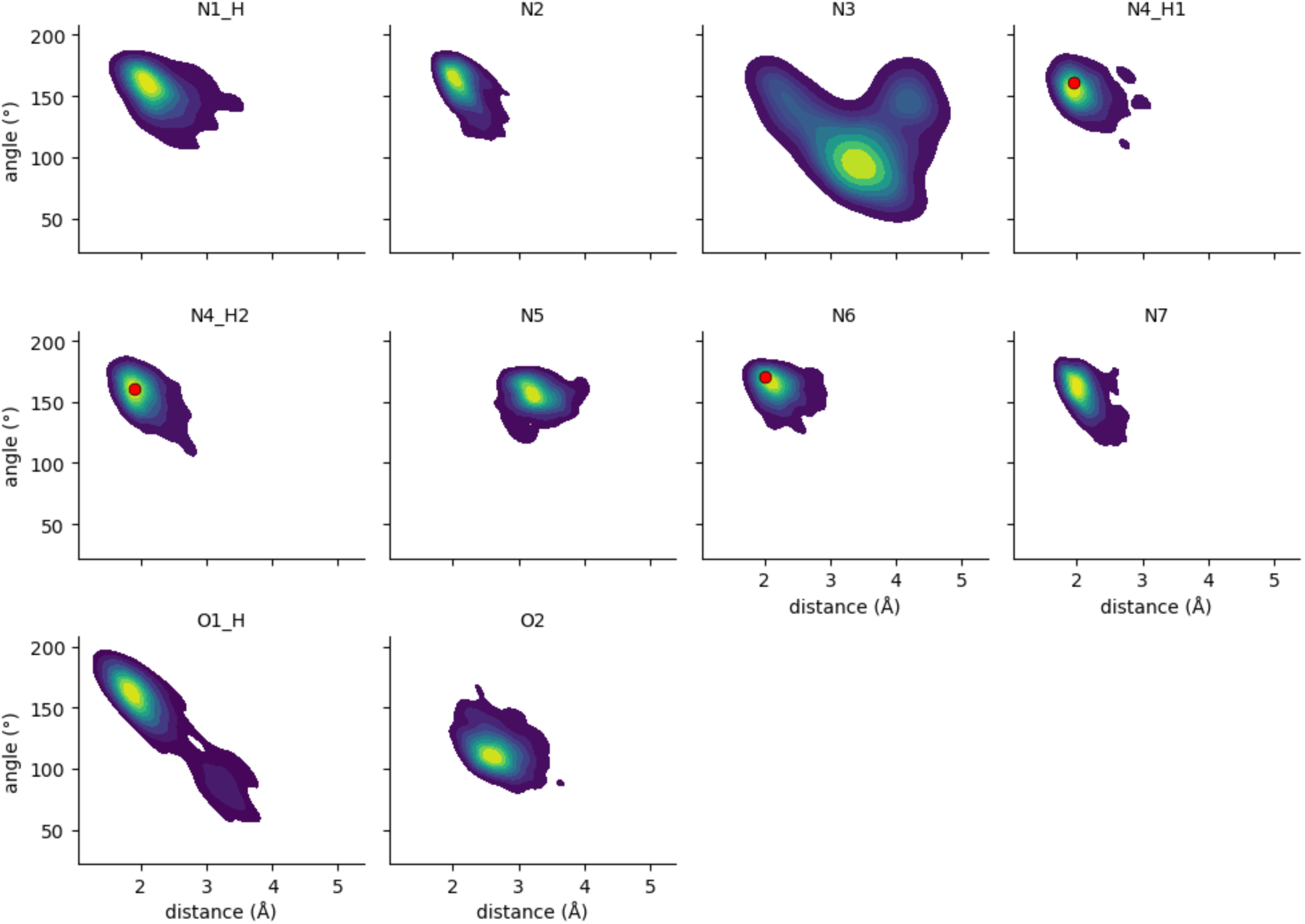
Hydrogen-bond geometry for each polar donor (D) / acceptor (A) atom of ZMA. The colored maps are joint distributions of the H···A distance (x-axis) and D–H···A angle (y-axis) in the bound state for the WT hA_2A_R. Red points indicate the hydrogen-bond geometry with the protein in the crystal structure of the WT hA_2A_R.

**Fig. S4.**
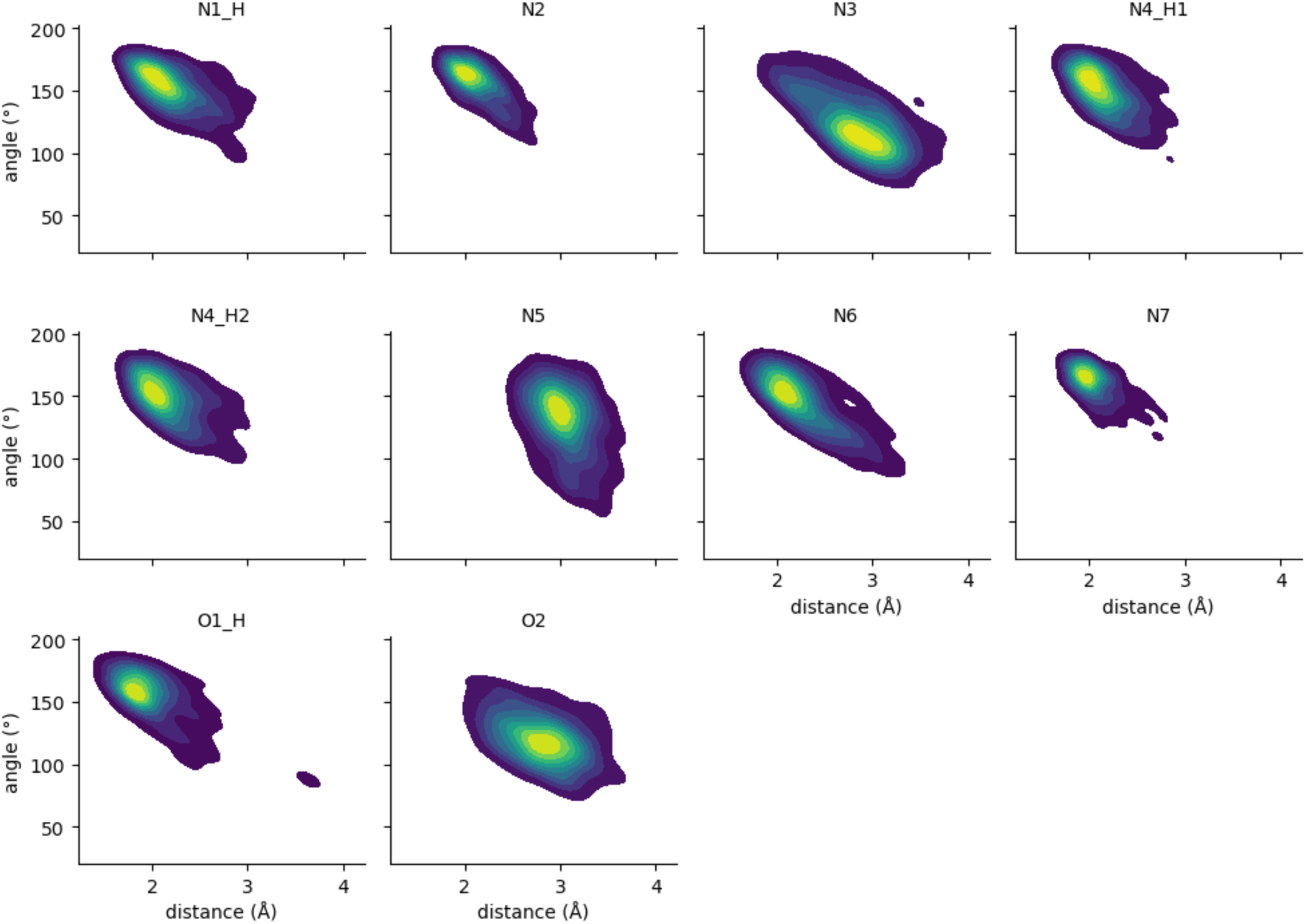
Hydrogen-bond geometry for each polar donor (D) / acceptor (A) atom of ZMA. The colored maps are joint distributions of the H···A distance (x-axis) and D–H···A angle (y-axis) in the TS for the WT hA_2A_R.

**Fig. S5.**
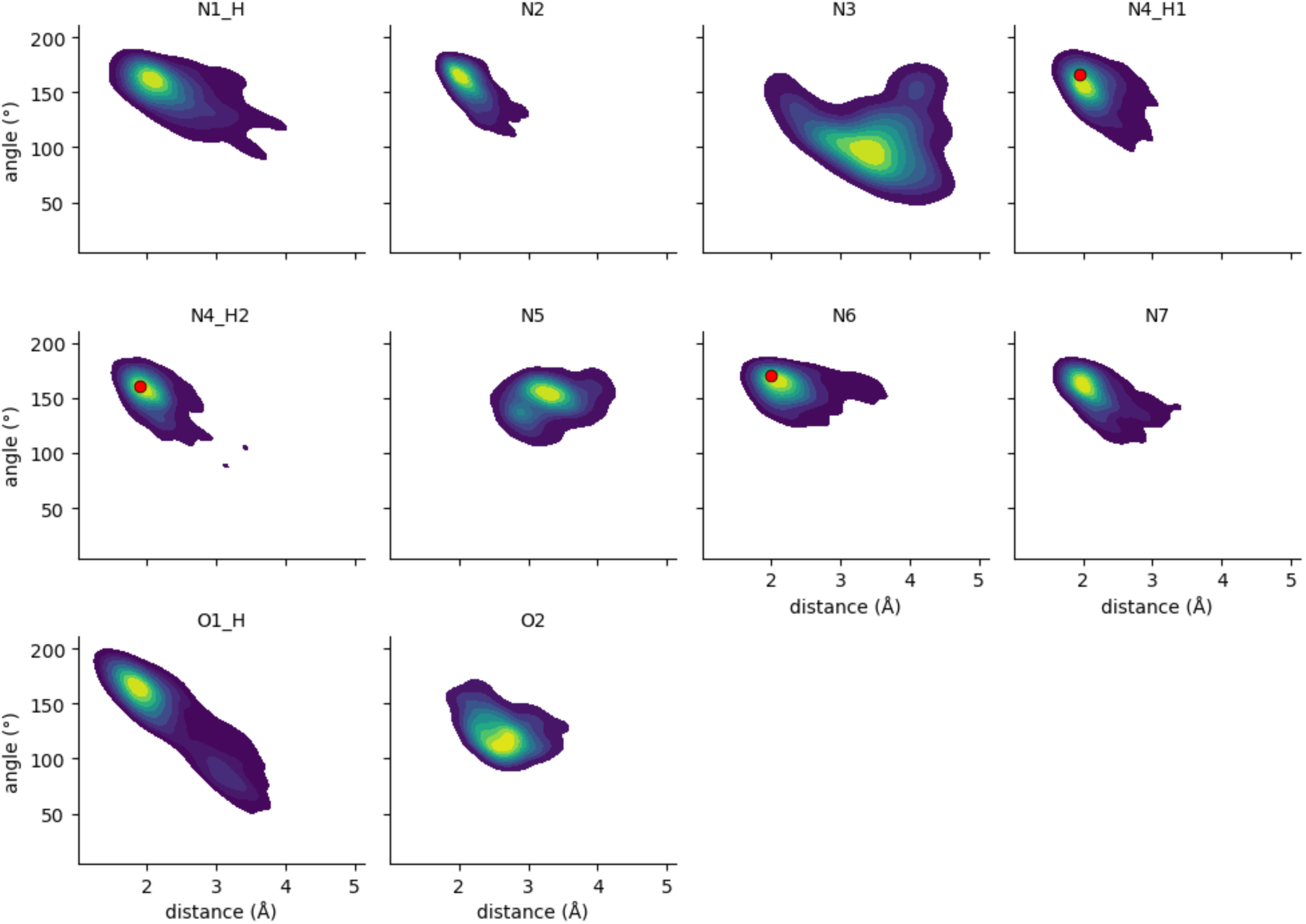
Hydrogen-bond geometry for each polar donor (D) / acceptor (A) atom of ZMA. The colored maps are joint distributions of the H···A distance (x-axis) and D–H···A angle (y-axis) in the bound state for the EtoQ mutant. Red points indicate the hydrogen-bond geometry with the protein in the crystal structure of the WT hA_2A_R.

**Fig. S6.**
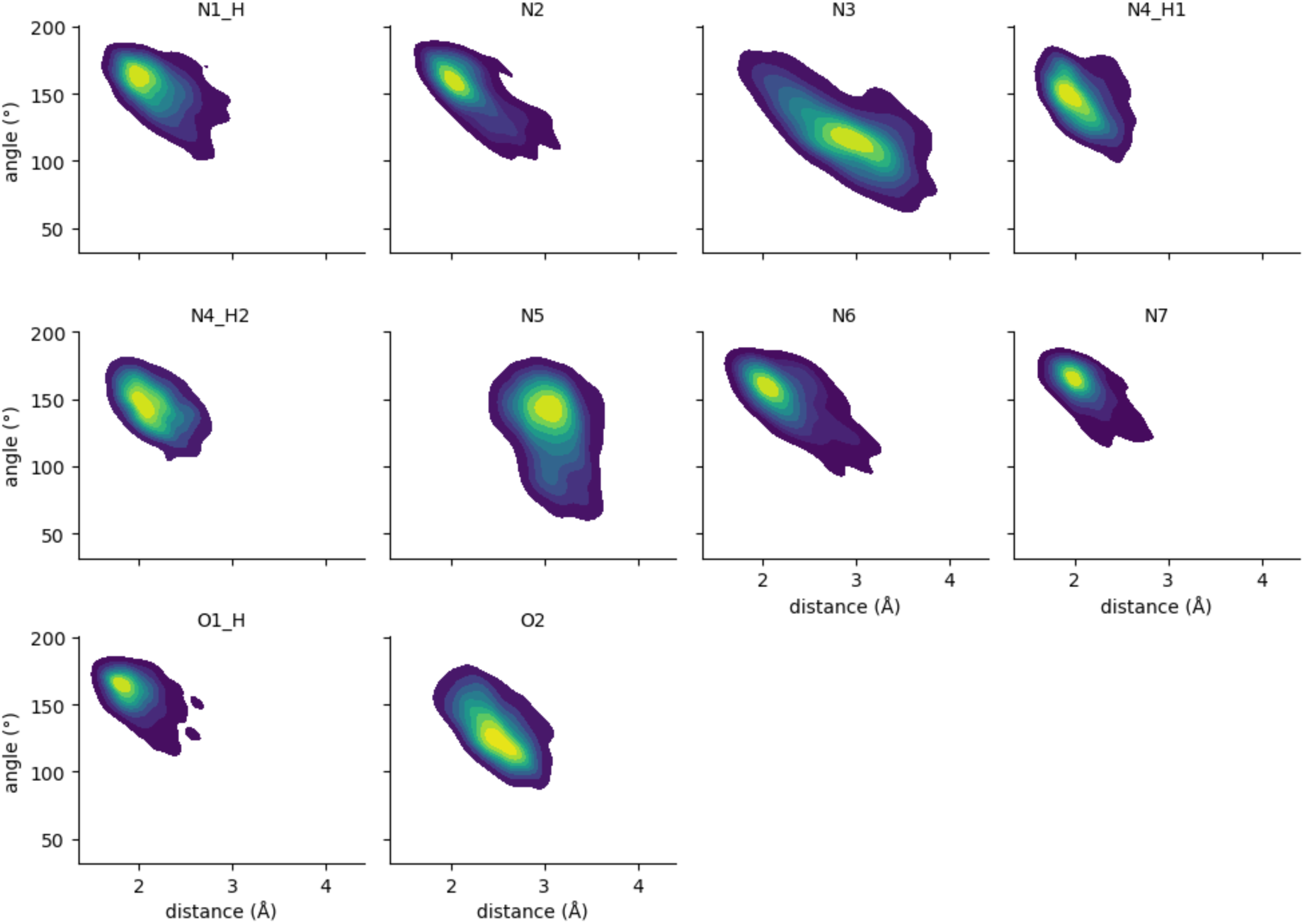
Hydrogen-bond geometry for each polar donor (D) / acceptor (A) atom of ZMA. The colored maps are joint distributions of the H···A distance (x-axis) and D–H···A angle (y-axis) in the TS for the EtoQ mutant.

**Fig. S7.**
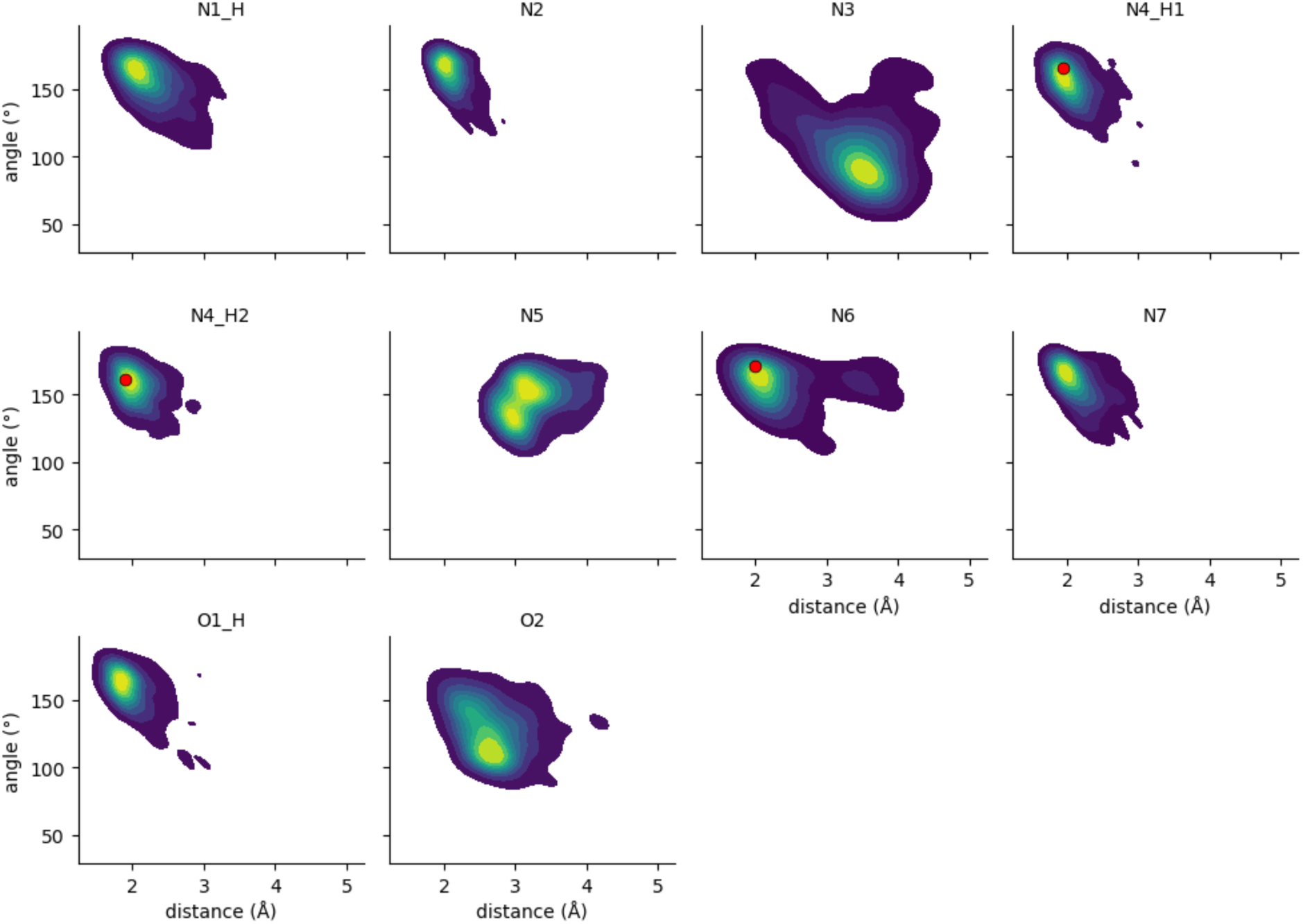
Hydrogen-bond geometry for each polar donor (D) / acceptor (A) atom of ZMA. The colored maps are joint distributions of the H···A distance (x-axis) and D–H···A angle (y-axis) in the bound state for the TtoA mutant. Red points indicate the hydrogen-bond geometry with the protein in the crystal structure of the WT hA_2A_R.

**Fig. S8.**
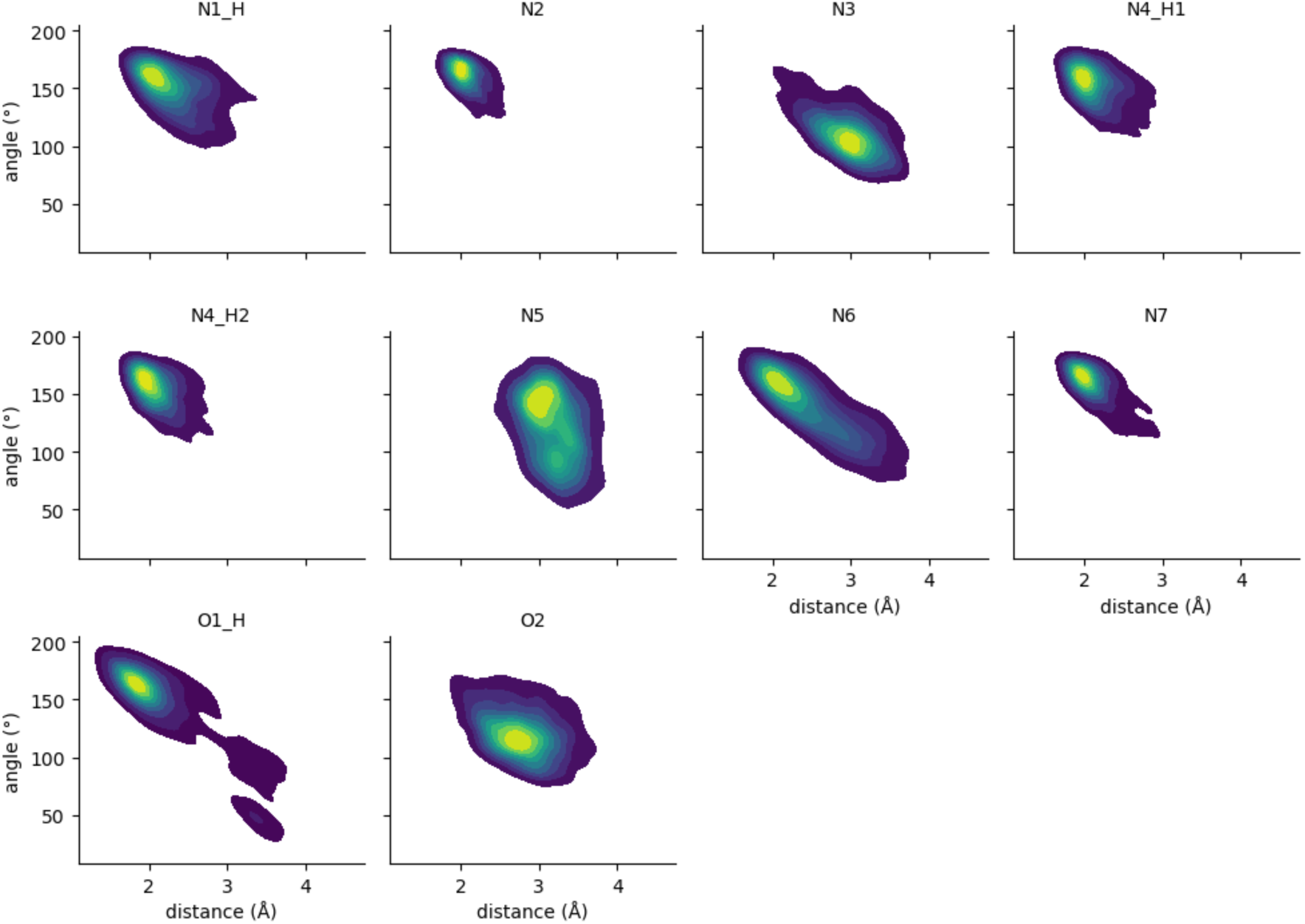
Hydrogen-bond geometry for each polar donor (D) / acceptor (A) atom of ZMA. The colored maps are joint distributions of the H···A distance (x-axis) and D–H···A angle (y-axis) in the TS for the TtoA mutant.

**Fig. S9.**
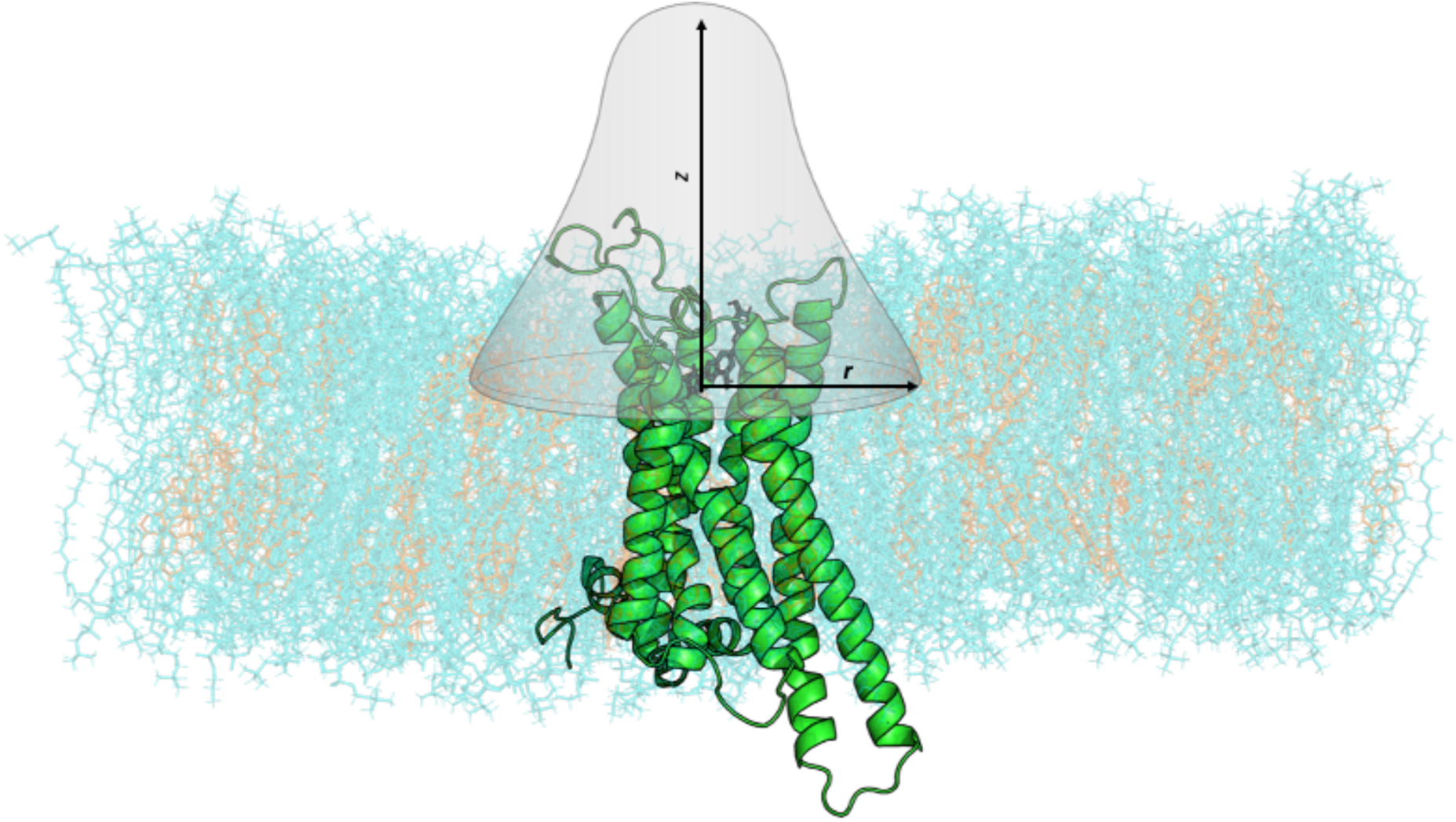
Schematic of the enhanced-sampling simulation setup, showing the restraining harmonic wall acting on the ligand center-of-mass (grey surface) and the definition of the two CVs *z* and *r*.

**Fig. S10.**
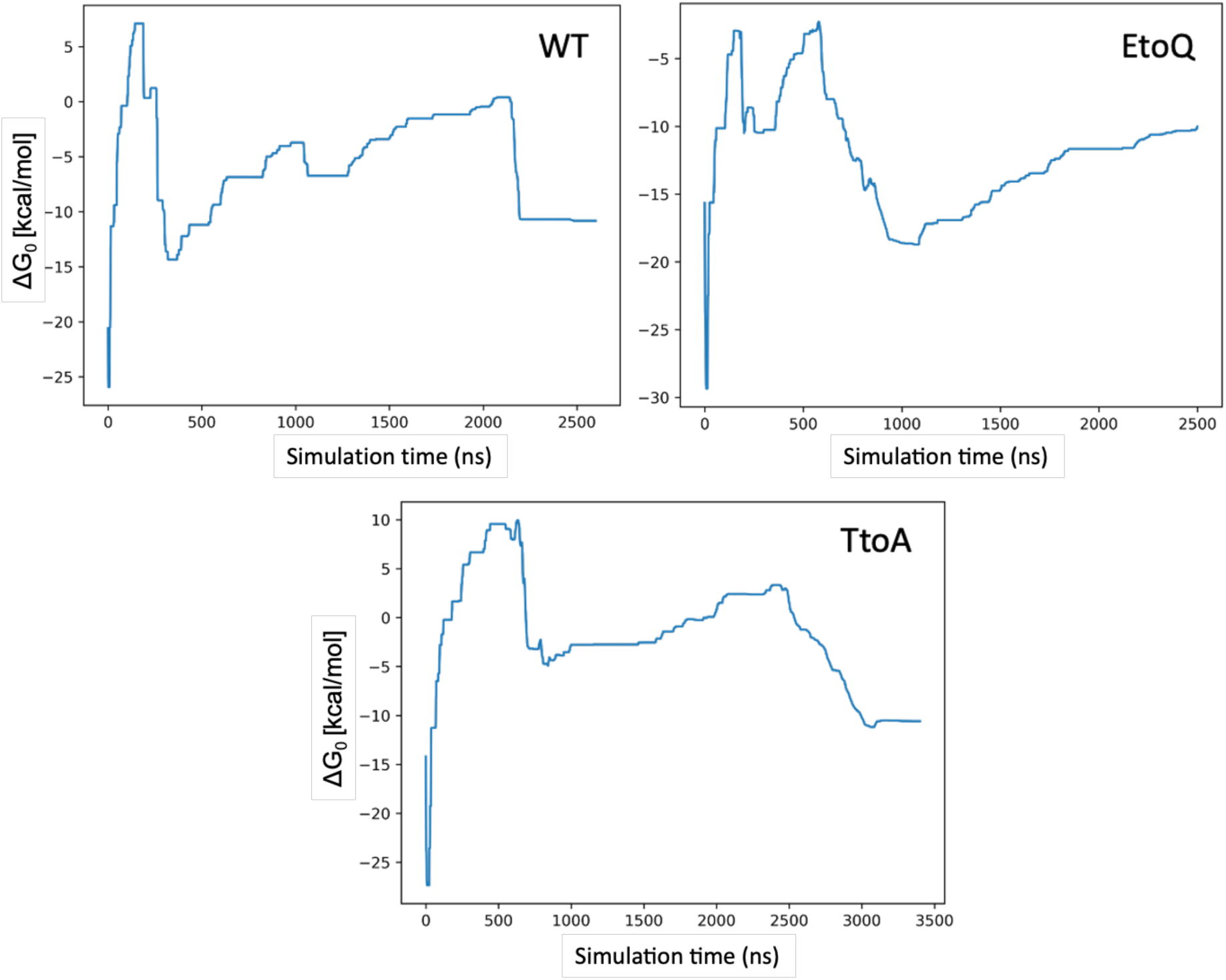
Time evolution of the volume-dependent binding free energy ΔG_0_ in the WT-MetaD simulations for WT hA_2A_R, and its EtoQ and TtoA mutants.

**Fig. S11.**
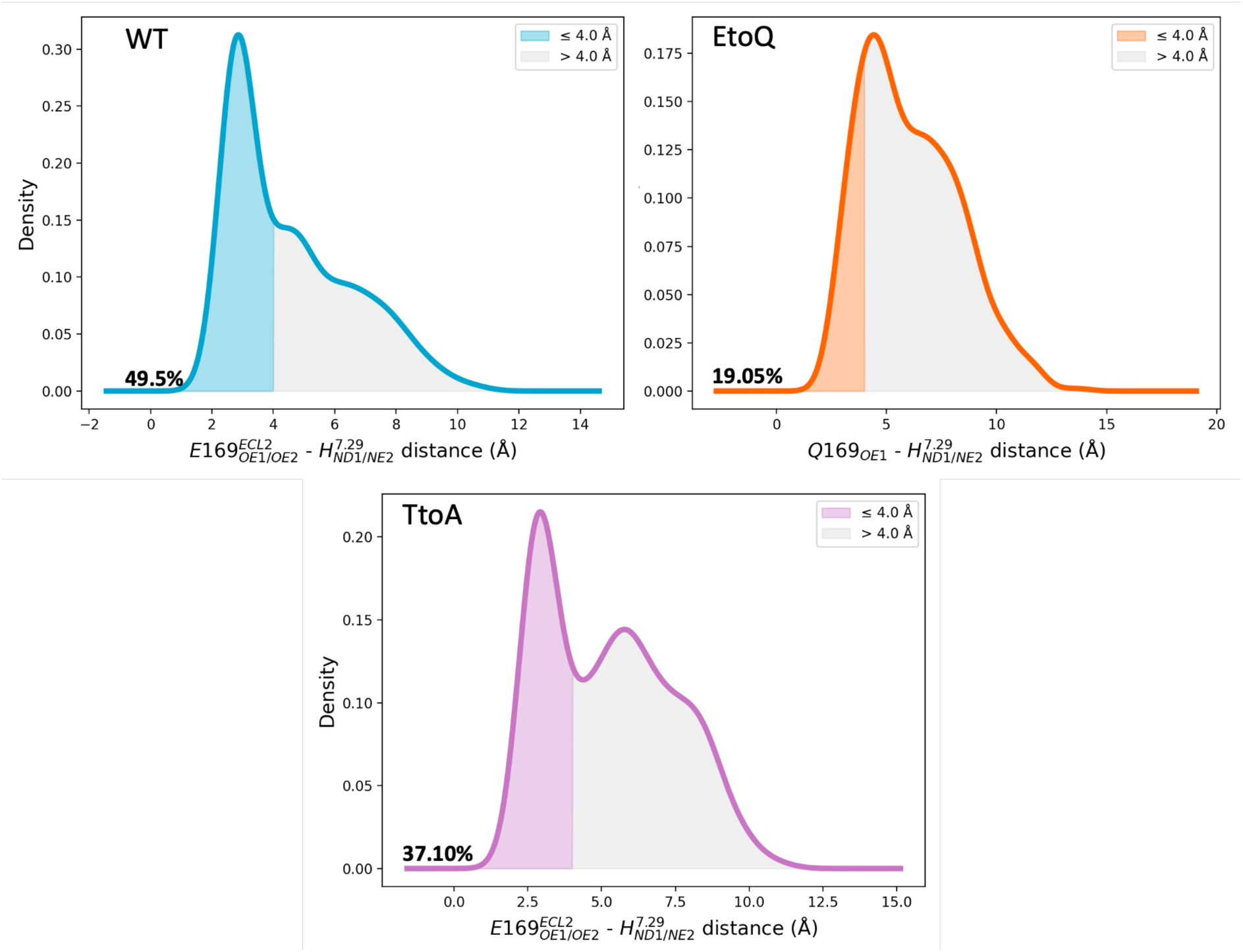
Bound state. Distributions of the salt bridge distance obtained in unbiased simulations of WT hA_2A_R and its EtoQ and TtoA mutants. The shaded regions denote configurations with (O-N) distances ≤ 4.0 Å (salt bridge formed), and the percentages indicate the corresponding population in each state. HtoA mutant is not shown, since here the salt bridge interaction is completely lost due to the mutation of the histidine in alanine.

**Table S1.** (Top) Bound state. Average number of water molecules within 0.35 nm of the heavy atoms of the ligand when bound or fully solvated. **(Bottom)** Average number of water molecules inside the binding pocket with the ligand bound or unbound. Results from WT-MetaD simulations. Uncertainties are SD.

|  | WT | EtoQ | TtoA |
| --- | --- | --- | --- |
| Ligand bound | $8 \pm 2$ | $8 \pm 2$ | $9 \pm 1$ |
| Ligand solvated | $29 \pm 4$ | $29 \pm 3$ | $31 \pm 4$ |
| Pocket (bound) | $30 \pm 4$ | $35 \pm 4$ | $36 \pm 5$ |
| Pocket (unbound) | $40 \pm 6$ | $47 \pm 5$ | $48 \pm 4$ |

**Table S2.** Bound state. Presence (**v**) / absence (**x**) of hydrophobic contacts between the ligand and the reported protein residues in the WT hA_2A_R, EtoQ and TtoA mutants. Results from WT-MetaD simulations. Uncertainties are SD. Presence was defined as an interaction occurring in >70% of the frames. The analysis was performed with the ProLIF software.

|  | I<sup>2.64</sup> | S<sup>2.65</sup> | V<sup>3.32</sup> | L<sup>3.33</sup> | M<sup>5.38</sup> | W<sup>6.48</sup> | L<sup>6.51</sup> | H<sup>6.52</sup> | L<sup>7.32</sup> | M<sup>7.35</sup> | Y<sup>7.36</sup> | I<sup>7.39</sup> |
| --- | --- | --- | --- | --- | --- | --- | --- | --- | --- | --- | --- | --- |
| WT | ✓ | ✓ | ✓ | ✓ | ✓ | ✓ | ✓ | ✓ | ✓ | ✓ | ✓ | ✓ |
| EtoQ | ✓ | ✓ | ✓ | ✓ | ✓ | ✓ | ✓ | ✓ | ✓ | ✓ | ✓ | ✓ |
| TtoA | ✓ | ✗ | ✓ | ✓ | ✓ | ✓ | ✓ | ✓ | ✓ | ✓ | ✓ | ✓ |

**Table S3.** Transition state. Average number of water molecules within different cutoff distances of the heavy atoms of the ligand. Results from WT-MetaD simulations. Uncertainties are SD.

| Cutoff (nm) | WT | EtoQ | TtoA |
| --- | --- | --- | --- |
| 0.35 | 21 ± 3 | 22 ± 3 | 20 ± 3 |
| 0.5 | 53 ± 4 | 58 ± 3 | 57 ± 3 |

## Footnotes

1 IUPAC name 4-(2-(7-amino-2-(furan-2-yl)-[1,2,4]triazolo[1,5-a][1,3,5]triazin-5-ylamino)ethyl)phenol.

2 The H-bond with E169^ECL2^ is often replaced by water, possibly because the mutation of T^6.58^ into alanine strongly destabilizes the interacting E169^ECL2^-T^6.58^-H^7.29^ triad in the WT (see SI Section 1).

